# An Ensemble Learning Approach to perform Link Prediction on Large Scale Biomedical Knowledge Graphs for Drug Repurposing and Discovery

**DOI:** 10.1101/2023.03.19.533306

**Authors:** Vignesh Prabhakar, Chau Vu, Jennifer Crawford, Joseph Waite, Kai Liu

**Affiliations:** Genentech Inc., 1 DNA Way, South San Francisco, CA - 94080, USA; Roche, 7070 Mississauga Rd, Mississauga, ON - L5N5M8, Canada

## Abstract

Generating knowledge graph embeddings (KGEs) to represent entities (nodes) and relations (edges) in large scale knowledge graph datasets has been a challenging problem in representation learning. This is primarily because the embeddings / vector representations that are required to encode the full scope of data in a large heterogeneous graph needs to have a high dimensionality. The orientation of a large number of vectors requires a lot of space which is achieved by projecting the embeddings to higher dimensions. This is not a scalable solution especially when we expect the knowledge graph to grow in size in order to incorporate more data. Any efforts to constrain the embeddings to lower number of dimensions could be problematic as insufficient space to spatially orient the large number of embeddings / vector representations within limited number of dimensions could lead to poor inferencing on downstream tasks such as link prediction which leverage these embeddings to predict the likelihood of existence of a link between two or more entities in a knowledge graph. This is especially the case with large biomedical knowledge graphs which relate several diverse entities such as genes, diseases, signaling pathways, biological functions etc. that are clinically relevant for the application of KGs to drug discovery. The size of the biomedical knowledge graphs are therefore much larger compared to typical benchmark knowledge graph datasets. This poses a huge challenge in generating embeddings / vector representations of good quality to represent the latent semantic structure of the graph. Attempts to circumvent this challenge by increasing the dimensionality of the embeddings often render hardware limitations as generating high dimensional embeddings is computationally expensive and often times infeasible. To practically deal with representing the latent structure of such large scale knowledge graphs (KGs), our work proposes an ensemble learning model in which the full knowledge graph is sampled into several smaller subgraphs and KGE models generate embeddings for each individual subgraph. The results of link prediction from the KGE models trained on each subgraph are then aggregated to generate a consolidated set of link predictions across the full knowledge graph. The experimental results demonstrated significant improvement in rank-based evaluation metrics on task specific link predictions as well as general link predictions on four open-sourced biomedical knowledge graph datasets.

## 1 Introduction

A knowledge graph (KG) is a collection of known facts in the form of a directed labeled heterogeneous graph, wherein each node represents an entity and each edge represents a relation between the entities. Each fact is represented as a triple of the form (head entity, relation, tail entity); for example, the fact that Berlin is the capital of Germany can be stored as the triple (‘Berlin’, ‘capital of’, ‘Germany’). The categories or classes of entities and relations in the knowledge graph are standardized to a closed set. Knowledge graphs were initially utilized for graph based structured data storage and retrieval of facts via computationally simple graph traversal algorithms but recently the task of predicting missing links between entities in the graph has become an active area of research due to the potential of identifying novel relationship between entities while predicting the missing links in the graph.

While many approaches for knowledge graph link prediction have been explored, knowledge graph embedding has gained much traction recently [1]. Knowledge graph embedding (KGE) models with an optimization strategy can generate embeddings / vector representations which capture latent properties of the entities and relations in the graph [2]. These embeddings can then be used in downstream machine learning tasks such as node classification [3], community detection [4], and in our case link prediction [1]. In general, the likelihood of existence of a missing link between two entities in the KG can be predicted by computing the proximity of a head entity embedding and relation embedding with that of a tail entity embedding by passing them as inputs to model-specific scoring functions as shown in Table 1 and computing aplausibility/confidence score for the existence of a link between the two entities [5].

**Table 1.** Scoring functions of standard knowledge graph embedding models given the head entity vector representation h, tail entity vector representation t and relation vector representation r

|  | Embedding Model | Score Function |
| --- | --- | --- |
| Translational Distance Models | TransE | $F(h, r, t) = -\ h + r - t\ $ |
| | ComplEx | $F(h, r, t) = Re(< h, r, t' >)$ |
| | RotatE | $F(h, r, t) = \ -h \cdot r - t\ $ |
| Semantic Matching Models | RESCAL | $F(h, r, t) = h^T W_r t$ |
| | DistMult | $F(h, r, t) = < h, r, t >$ |

Even though these embedding models have demonstrated robust performance on benchmark datasets [6–8]; previous work has shown that biomedical knowledge graphs have a distinct topological structure [9], are very large in size and incorporate various categories of information in their nodes and edges compared to typical benchmark graphs such as Freebase FB15k and Wordnet WN18 that have about 600,000 and 150,000 triples respectively and incorporate fewer categories of nodes and edges to represent heterogeneous information. Smaller biomedical knowledge graph datasets such as BioKG and Hetionet have approximately 2M and 2.2M triples respectively [10–12] and larger biomedical knowledge graphs such as Drug repurposing knowledge graph ( DRKG ) and Integrated biomedical knowledge hub ( iBKH ) have approximately 6M and 50M triplets respectively [13, 14]. Datasets of this size could still be tenable for training, given a high-performance computing infrastructure with multiple GPUs. However, as additional triples are added to further augment the knowledge base, these graphs may grow too enormous. Therefore the task of generating low dimensional robust representations for the entities and relations in such a knowledge graph becomes complicated as the model can only incorporate a limited number of entity and relation embeddings within a given number of dimensions while still avoiding congestion of the embedding vectors and maintaining the semantic meaning conveyed by the graph. Moreover, tackling this problem by allocating more computational resources is neither acost-effective nor scalable solution.

In order to generate embeddings / vector representations for the entities and relations in a large knowledge graph while ensuring that the embeddings effectively represent the latent semantic properties of the graph in lower dimensions; this paper proposes an ensemble approach for performing KGE model training on subgraphs (localized view of full data) sampled from the full graph and aggregating plausibility scores for link predictions obtained from each subgraph to generate a consolidated set of link predictions across the full KG.

More specifically, in order to avoid the very high dimensional embeddings required for representing latent properties of a large scale knowledge graph, we propose a stochastic sampling approach in which few random seed nodes are chosen first, then breadth-first search (BFS) is performed to add other nodes and edges that are connected to the chosen seed nodes to the subgraph until each generated subgraph reaches a specific size that is determined by the number of triples contained in the subgraph [15]. Each of these subgraphs can be considered as a localized view space of the full scope of the KG. KGE models are trained individually on each subgraph wherein each triple (*h, r, t*) is assigned a plausibility/confidence score based on the scoring function of the corresponding subgraph’s embedding model. The ranked-triples /link predictions along with their respective plausibility scores from each subgraph are then aggregated to yield a consolidated set of ranked-triples / link predictions across the full KG. In order to generate the ranking for a triple/ link-prediction in the context of the full KG using the ensemble learner; for a particular triple *T* that exists in multiple subgraphs the plausibility scores for all the triples in those individual subgraphs are firstly normalized before aggregating the plausibility score for triple *T* across all those subgraphs using a simple average of it’s normalized plausibility scores from every subgraph in which it is present. The averaged plausibility score is considered the final score assigned to the triple *T* in the full KG. Similarly aggregated plausibility scores are computed for the remaining triples for generating the consolidated set of ranked-triples / link predictions across the full KG.

Experimental results generated on typical benchmark biomedical KG datasets showed that our ensemble approach yielded significantly better performance for link prediction on large biomedical KGs while consuming lesser GPU memory footprint compared to the traditional approach of training a KGE model on the full knowledge graph.

## 2 Literature Review

Knowledge graph embedding models represent entities and relations as low-dimensional vectors to use in downstream applications, including node classification and link prediction. Specifically, link prediction task leverages these latent representations as inputs to a scoring function to compute the plausibility of a link existing between a particular pair of nodes/entities in the KG. Various scoring functions have been proposed in recent years [16], including distance-based scoring functions used by translational distance models such as TransE, ComplEx, RotatE etc. [7, 17, 18] and similarity-based scoring functions used by semantic-matching models such as RECSAL, DistMult etc. [19, 20]. Below in Table.1, we list a few well-known knowledge graph embedding models and their corresponding scoring functions [21].

In Table.1; the function *F* (*h, r, t*) represents the scoring function of the embedding model that is used to rank the likelihood of the existence of a link between a head-tail entity pair. *h* represents the vector representation of the head entity, *t* represents the vector representation of the tail entity and *r* represents the vector representation of relation connecting the head entity h with the tail entity *t*.

There have been some previous studies that have explored the possibility of generating vector representations for a local view space (subgraphs) instead of a global view space (full KG) so as to circumvent the challenges arising due to the high dimensionality of embeddings required to represent entities and relations in the global view space. The primary idea behind doing this has been that the full scope of the KG is not required to generate vector representations / embeddings for any given node/edge in the graph and only the nodes and edges present in the neighborhood of a particular node or edge impact it’s embeddings. Therefore; any kind of downstream inferencing will also require only a small subset of nodes and edges that lie within the neighborhood of a particular node/edge involved in the inferencing task. Some of these approaches include inductive embeddings, graph embeddings, utilizing non-euclidean embedding spaces and ensembling multiple embedding spaces.

Inductive Embeddings Approach: Apart from graph compression approaches [22], few recent works have explored avenues for training large-scale knowledge graphs using inductive embedding models. The knowledge graph embedding models mentioned in Table 1. are transductive embedding models which require full scope of the KG in order to generate vector representations for the nodes and edges in the KG. On the other hand, inductive embedding models create vector representations for any node or edge in the knowledge graph using just a few nodes and edges in the knowledge graph which are in the neighborhood of a given node or edge for which the vector representations are already computed. Several inductive embedding techniques have been proposed to help generate scalable vector representations on large graphs [23, 24]. Notably, Hamilton et al. introduced an inductive framework that leverages a graph convolutional network and learns feature aggregator functions to generate node embeddings [25]. Since this framework can generalize to unseen nodes for a large knowledge graph, this method can theoretically be trained using a smaller subgraph and applied on the remainder of the full graph for subsequent embedding generation and link prediction. However, this framework assumes that the unseen nodes are surrounded by known entities/nodes. Therefore; the training subgraph still needs to be of sufficient size to be able to produce robust results.

Graph embeddings approach: To address training on very large-scale graphs, a divide-and-conquer approach can be applied. One possible direction that has been explored previously involves dividing the full knowledge graph into smaller subgraphs and performing inferencing on each subgraph. However, this requires an aggregation scheme to combine results from each subgraph and consolidate. Graph embedding methods that encode individual subgraphs in entirety (instead of encoding nodes and edges within the graphs) as low-dimensional vectors offers a potential solution [26, 27], where these subgraph vector representations can then be fed into a classification framework to determine whether each subgraph is suitable to perform a particular link prediction task. Teru et al. proposed a task-specific inferencing approach in which a subgraph that encloses the specific target entities is extracted from the full knowledge graph and a graph neural network is used to score the likelihood of a specific triple given the extracted structure of the subgraph [28]. Another graph neural network (GNN) based approach is to train the GNN using subgraphs as mini-batches instead of training a GNN on the full original graph. This method achieved robust performance with less training time and memory resource requirements [29].

Ensemble approach: Previous publications have worked with ensemble methods to reduce variance and improve performance on knowledge graph completion. Wan et al. proposed an ensemble approach to extract multiple subgraphs of approximately 70% the size of the original knowledge graphs and perform representation learning on each subgraph before applying adaptive averaging on the entity embeddings in order to denoise embedding models for the full graph [30]. Despite the improvement in evaluation metrics, this method did not consider cases in which the learned node embeddings are vastly different between subgraphs (due to low overlap between subgraphs or different random seeds) and embedding vectors cannot be averaged. On the other hand, instead of averaging the embedding vectors, Xu et al. performed ensemble aggregation by averaging the scores for each triple calculated from different score functions for each embedding model ( Table 1) [31]. Similarly, Tay et al. introduced puTransE which trains on multiple embedding spaces, each with constrained selection of triplets from the original graph; the final score of each triple is the maximum TransE score function out of all subgraphs [32]. However, these approaches do not consider the case in which the score distributions for each subgraph training can be vastly different in ranges and scales. To correct this, Krompass et al. proposed score standardization to a [0, 1] range using a Platt scaler to demonstrate better performance using an ensemble model compared to an individual model [33].

Non-euclidean embedding spaces: The traditional knowledge graph embedding models operate in the euclidean space. Euclidean manifolds have zero curvature and therefore the amount of space to represent data in any dimension is limited. Therefore we often end up with high dimensionality when trying to get vector representations for large amounts of data in the euclidean space. However, Reimmanian manifolds have been explored before to represent large amounts of data in lesser number of dimensions [34, 35]. Hyperbolic manifold is one such example of a Reimmanian manifold. Hyperbolic manifolds have constant negative curvature and can contain a large number of vector representations in any given dimension. Therefore large amounts of data can be represented in lesser dimensions. This is simply done by projecting vectors from euclidean space to hyperbolic space by applying a hyperbolic norm to the vectors. The poincare ball model is one such model proposed by Maximilian Nickel and Douwe Kiela that creates vector representations / embeddings of the nodes and edges from a large scale graph in the hyperbolic space [36, 37]. This model is successful in creating low dimensional embeddings for large scale graph datasets but is not inductive in nature and cannot be applied to unseen nodes as the graph grows in size.

## 3 Methodology

We have leveraged an ensemble learning approach that aggregates link predictions generated from embeddings trained on multiple localized view spaces (subgraphs) to come up with a consolidated set of link predictions for the full KG. This is a divide and conquer algorithm where we divide the big KG into multiple smaller subgraphs of equivalent size and then train KGE models on those individual subgraphs to generate relevant ranked triples / link predictions per subgraph which is then combined using an aggregation strategy to produce link predictions for the full KG.

The broader overview of the steps that we have adopted in our approach have been depicted in Figure 1 as well as listed below:

**Figure 1.**
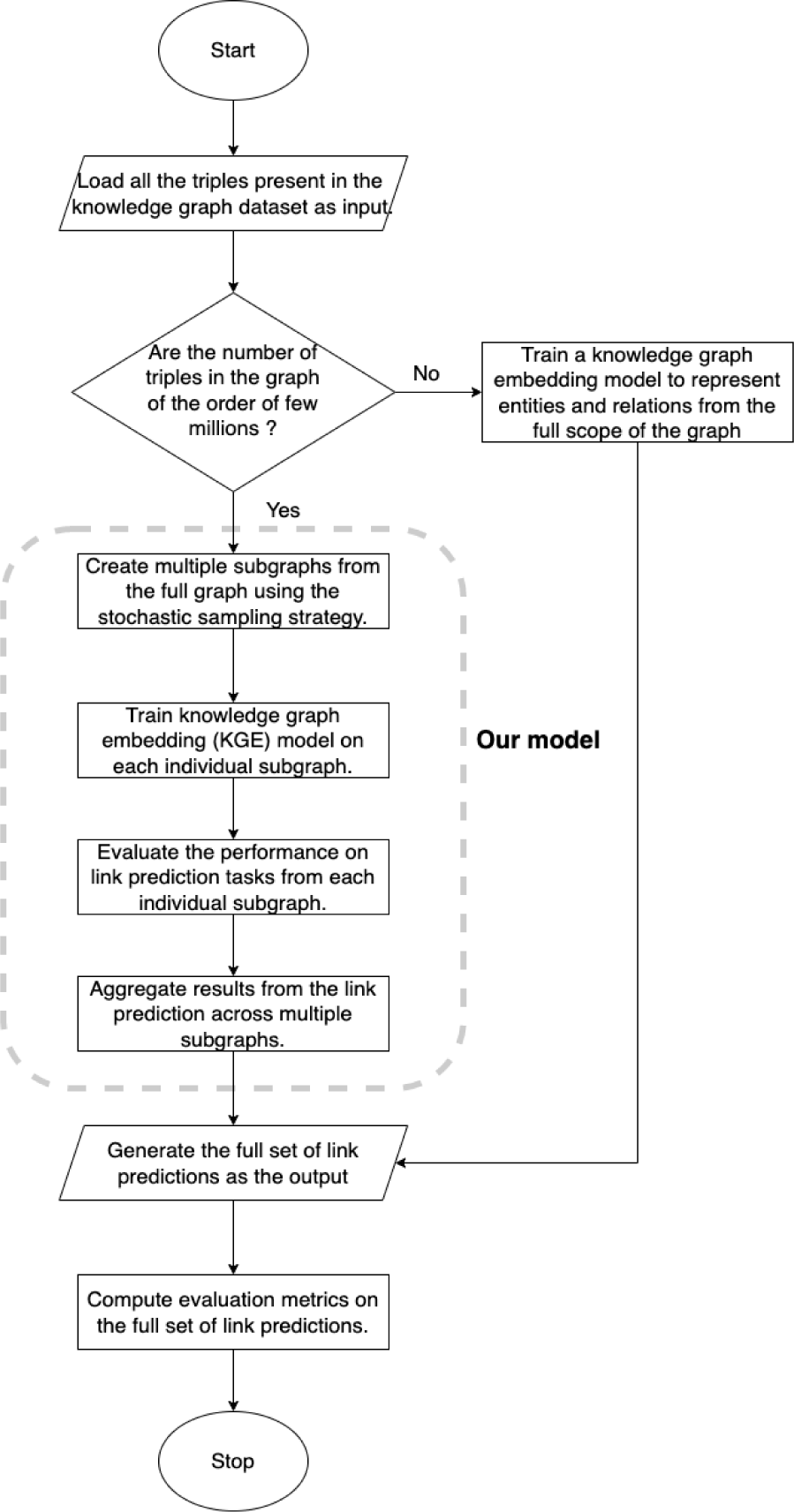
Flowchart showcasing the broader level overview of the steps involved in data preprocessing, training and evaluation of our ensemble learner.

1. Creating multiple subgraphs from the full knowledge graph dataset.
2. Train knowledge graph embedding model on each individual view / subgraph.
3. Evaluate the performance on link prediction tasks from each individual subgraph.
4. Aggregate the link predictions results from multiple subgraphs.

### 3.1 Algorithmic description

The inputs that are required for executing our algorithm are the full scope of the knowledge graph *G*, number of subgraphs to be sampled from *G* and the upper, lower bounds for the number of triples that can be present per sampled subgraph. Our algorithm scales well for large knowledge graphs and therefore we recommend having several million triples in the full KG so as to distinguish and compare the results of our approach shown in Algorithm 1 and Figure 4. against the traditional approach of training KGEs on full graph *G* shown in Figure 2. The primary steps involved in this algorithm are to select a random seed triple *T_j_* from the full knowledge graph *G* to get started with and then continuously add neighboring triples to this seed triple based on the adjacency matrix of *G* (using BFS) while ensuring that the subgraph being generated obeys the upper and lower bound constraints for the number of triples that can exist per subgraph. This is done to generate subgraphs of equivalent sizes. Once the bound limits are reached; we finish the process of sampling one subgraph as shown in Figure 3. Thereafter we train embedding models to generate vector representations for the entities and relations within the sampled subgraph and perform link prediction using the embeddings of the subgraph.

**Figure 2.**
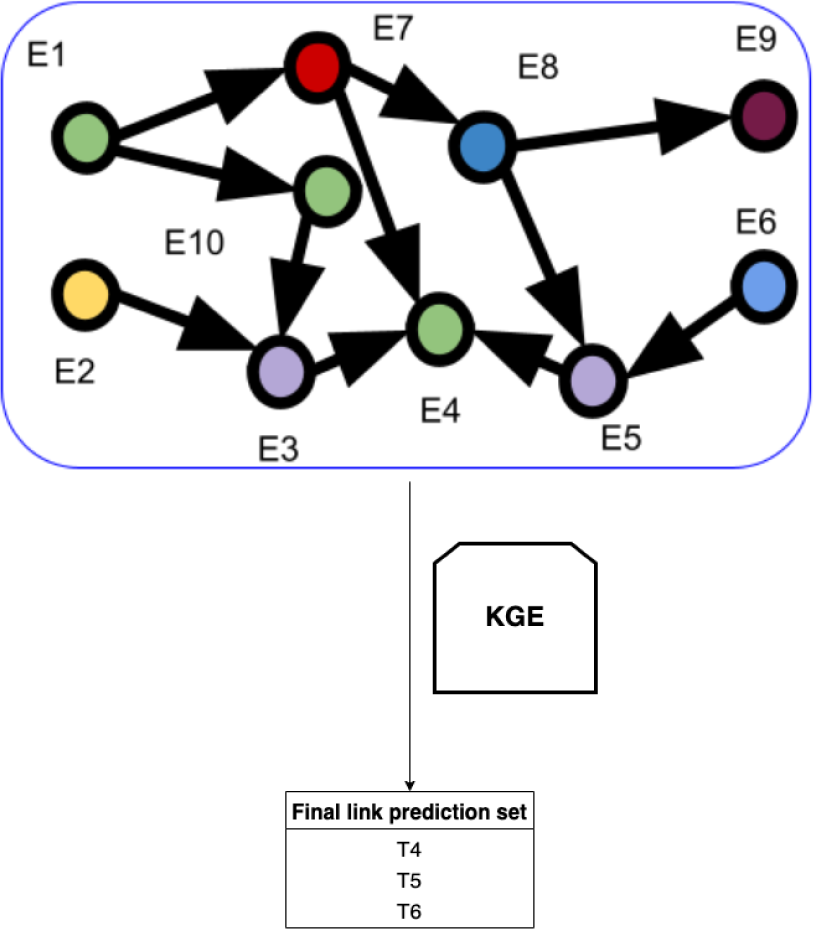
Architecture diagram of the traditional approach of training a knowledge graph embedding model on the full knowledge graph to generate link predictions. Here, *E*_1_*, E*_2_*, E*_3_etc. are the entities of the knowledge graph. *T*_4_*, T*_5_*, T*_6_ etc. represent the tail entities with the highest plausibility scores corresponding to the likelihood of existence of a link with agiven head entity.

**Figure 3.**
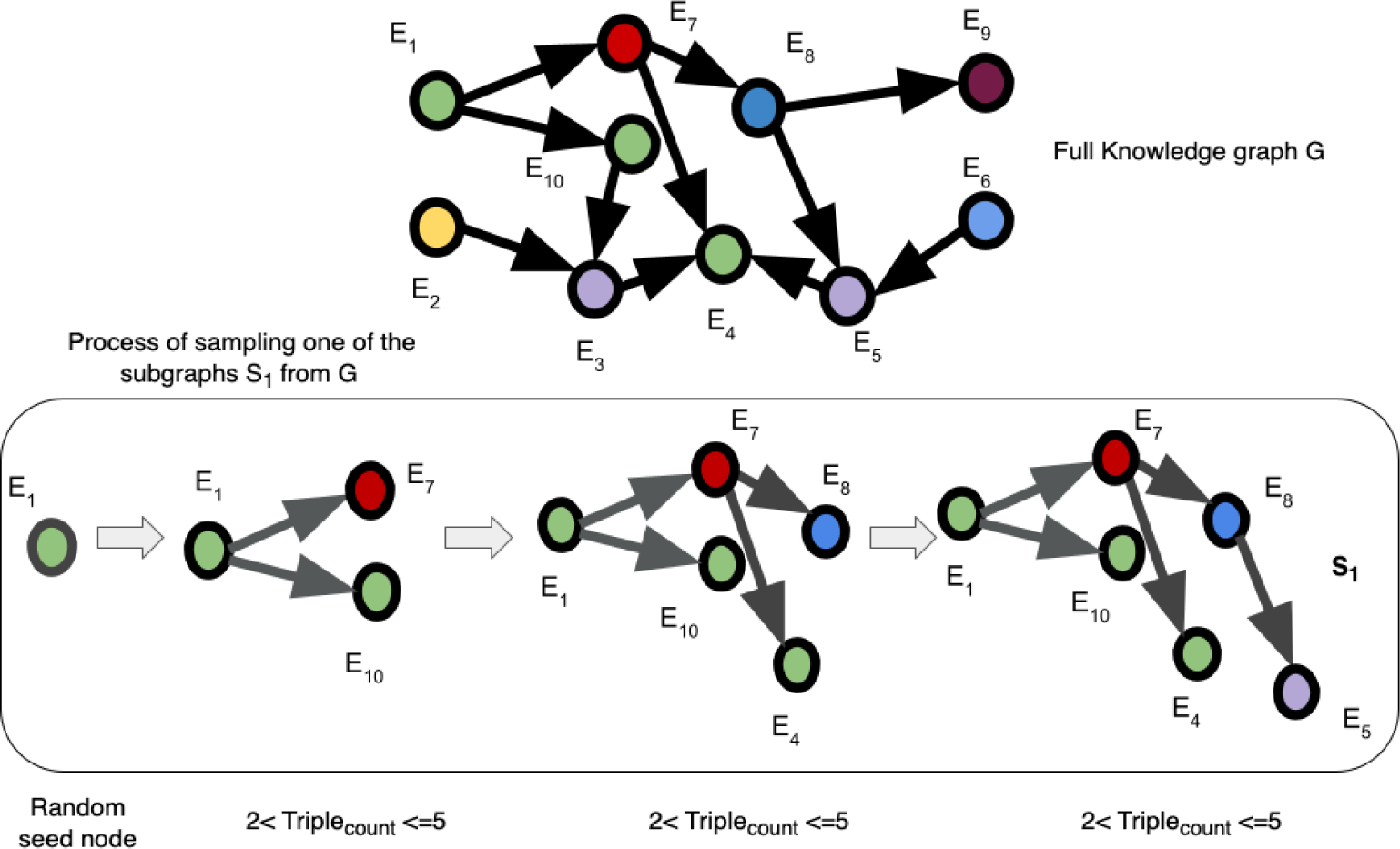
Illustration of the stochastic sampling approach adopted to create a single subgraph *S*_1_ from a full knowledge graph *G*. *E*_1_*, E*_2_*, E*_3_ etc. are the entities of *G*. We start with a random seed entity *E*_1_ and then add all the neighboring triples of the entities which are connected to our seed entity from the full knowledge graph *G* until we reach the specified bounds and only keep the largest connected component of the network to create the resultant subgraph *S*_1_.

**Figure 4.**
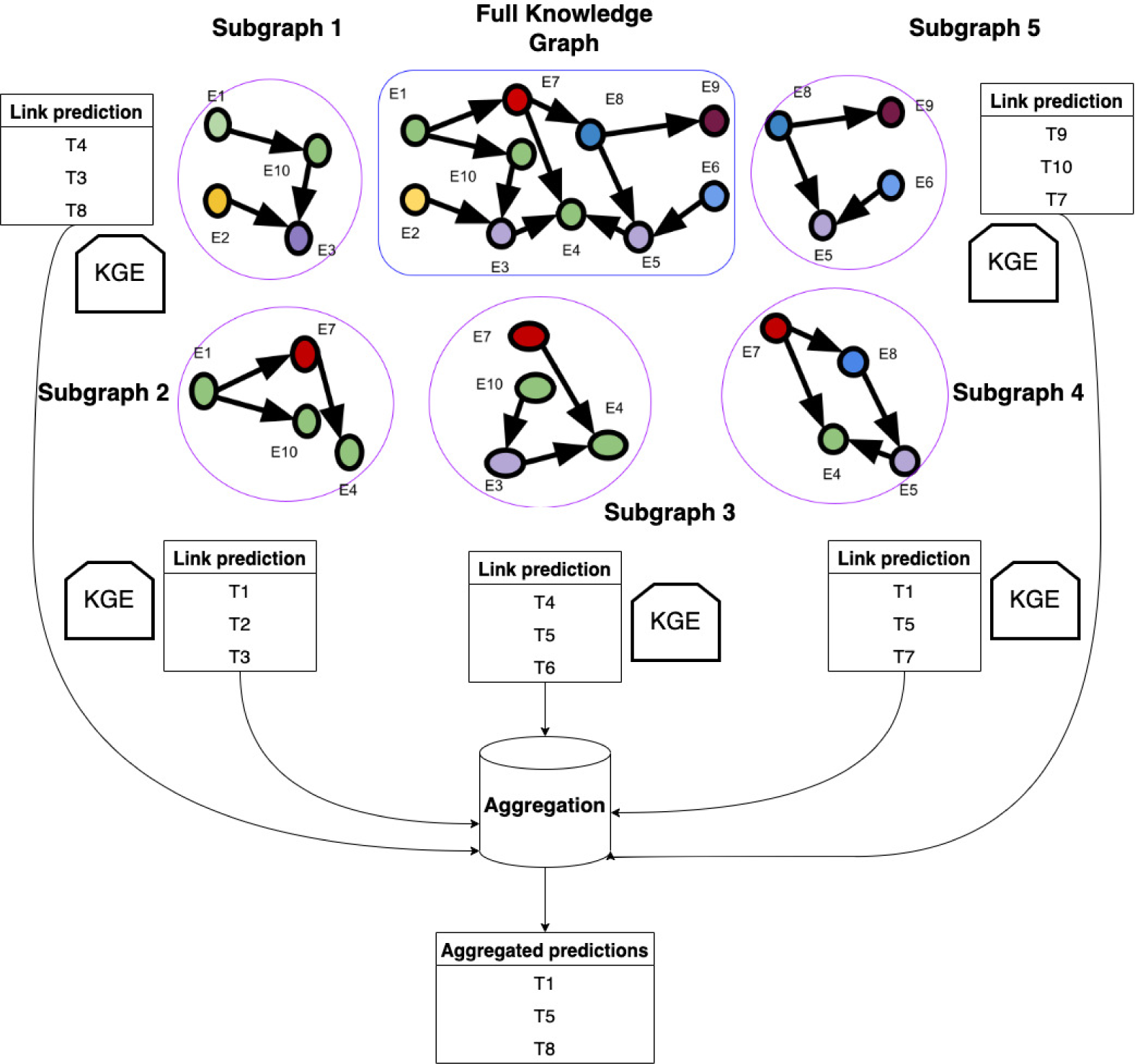
Architecture diagram of our Ensemble Learner for obtaining link predictions on large scale knowledge graphs. Here, *E*_1_*, E*_2_*, E*_3_ etc. are the entities of the large scale knowledge graph. *T*_1_*, T*_2_*, T*_3_*, T*_4_ etc. represent the tail entities with the highest plausibility scores corresponding to the likelihood of existence of a link with a given head entity in the knowledge graph.

The same process is iteratively repeated for generating remaining subgraphs from *G*, training embedding models on the subgraphs and consequently generating separate sets of link predictions from each of the subgraph.

These link predictions are generated along with their respective plausibility scores based on the scoring function adopted by the embedding model as given in Table 1. This plausibility score is used for creating an aggregated set of link predictions across the full graph. The plausibilty scores for the link predictions are normalized for each subgraph and then aggregated by applying either *median*(), *max*() or *mean*() operation over the normalized plausibility scores for generating the link predictions across the full graph. Finally we output the aggregated set of link predictions by sorting them based on the aggregated plausibility scores.

### 3.2 Creating multiple subgraphs from the knowledge graph

The subgraph sampling strategy aims at creating clusters of connected nodes / subgraphs from the complete knowledge graph. In order to ensure subgraph sizes are equivalent, we specify the upper and lower bounds for the total number of triples that could exist within each subgraph. For a general link prediction task, the strategy for creating each subgraph is as follows:

Let us start with a random seed triple (*h, r, t*) where *h* is the head entity, *t* is the tail entity and *r* is the relation connecting them together. The nodes and edges in the seed triple are the first components that are added while creating a subgraph. Thereafter; neighboring nodes and edges of *h* and *t* are continuously added to the subgraph using BFS until the number of triples in the subgraph is within the size limits specified by the upper and lower bounds. Addition of neighbors is done in batches to improve efficiency of the sampling process. Finally, only the largest connected component is kept and an individual subgraph is hence sampled as shown in Figure 3. The same process is repeated to sample the remaining subgraphs.

**Algorithm 1.**
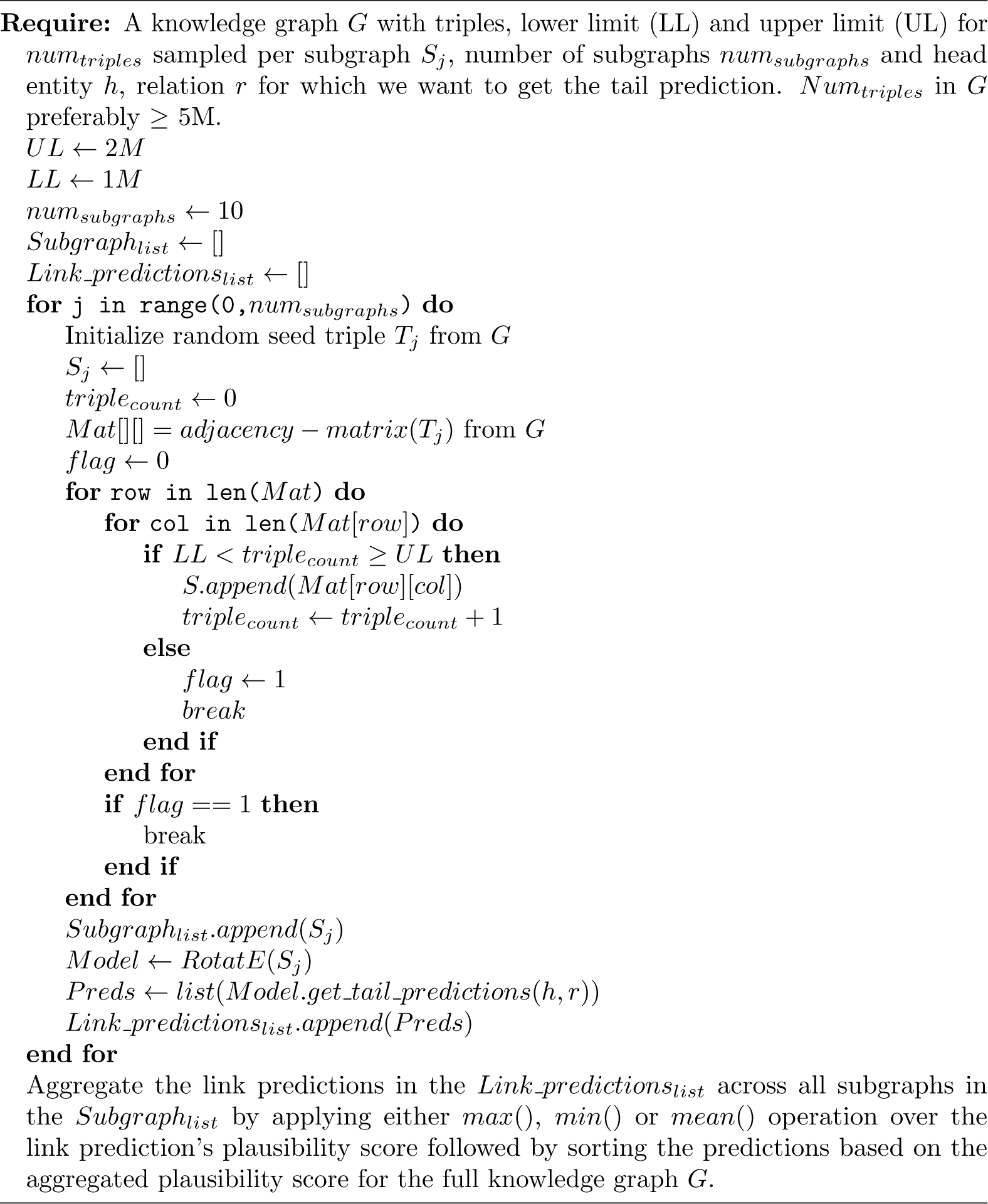
Ensemble learning for link prediction on large scale knowledge graphs – A divide and conquer approach:

A lot of times, knowledge graph completion aims at predicting links for a specific task, (e.g) a drug repurposing task requires predicting novel links between drug and disease entities in the graph. To incorporate such task-specific needs, the subgraph creation process is slightly modified. For a specific link prediction task between the head entity *E*_1_ and the tail entity *E*_2_, all entities of the same type/category as *E*_1_ and *E*_2_ are also added into a subgraph after the subgraph has been sampled by the process explained in Figure 3. This strategy ensures that each subgraph contains all the necessary entities that are required by the model for a specific link prediction task. Due diligence was taken to ensure that the final subgraph size still remained within the limits specified by the upper and lower bounds after adding all the task-specific entities that belong to the same category as *E*_1_ and *E*_2_.

### 3.3 Training knowledge graph embedding models on each individual subgraph

After generating all subgraphs; the next step is to train KGE models on each individual subgraph and generate embeddings for multiple local view spaces. These embeddings are then leveraged by the KGE’s scoring function to obtain link predictions from individual subgraphs. For the individual subgraph learners we use traditional knowledge graph embedding models listed in Table 1.

Some of the crucial hyperparameters that we optimize here are the number of sampled subgraphs, size of each sampled subgraph ( i.e the number of triples per subgraph ), embedding dimension size for KGEs trained on each subgraph, number of negative triples per positive triple in each subgraph, the margin for Margin Ranking Loss (also known as pairwise hinge loss) which is the loss function used by the individual subgraph learners and lastly the regularization coefficient.

Margin Ranking Loss is a linear learning-to-rank loss which is utilized for maximum-margin classification and can be used in pairwise settings to distinguish between the positive triples *T* ^+^ and negative triples *T ^−^* in the subgraphs with the goal being to maximize the difference between the plausibility scores assigned to *T* ^+^ and *T ^−^* by a good margin *λ* [38, 39]. The margin ranking loss / pairwise hinge loss is given by:

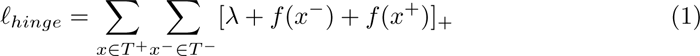

In equation 1 the objective is to optimize the loss function *ℓ_hinge_* towards generating embeddings that satisfies the condition *_x∈T_*+ *_x_−_∈T_− f* (*x*) *> f* (*x^−^*). Here, *f* (*h, r, t*) is the scoring function of the KGE used for generating embeddings for each subgraph, *x* is an instance of a positive triple *T* ^+^, *x^−^* is an instance of a negative triple *T ^−^* and *λ* is the margin used to maximize the difference between the plausibility scores assigned to *T* ^+^ and *T ^−^* which has to be selected by fine-tuning.

The margin for the loss function, number of negative triples per positive triple, embedding dimension size and regularization coefficient are hyperparameters that influence the optimization of the individual subgraph learners. The other hyperparamters such as number of sampled subgraphs and size of each sampled subgraph influence the optimization of the ensemble learner which is the net effect of combining link predictions from individual subgraph learners.

Apart from the hyperparameters specific to individual subgraph learners such as margin for the loss function; the other hyperparameters that are specific to the ensemble learner also need to be selected carefully by fine-tuning as they have significant impact on the performance of our ensemble learner. For example; a very large subgraph size leads to poor generalizability of the embeddings to the downstream link prediction task. This is because the entity and relation embeddings get affected by a lot of excessive information that serves as noise while computing the embeddings for a given enity/relation in the KG. A very small subgraph size leads to the generation of substandard embeddings due to insufficient data in the graph to create good vector representations for the entities and relations in the graph. Therefore, size of individual subgraphs is one of the crucial hyperparameters for our model that needs to be selected carefully.

Number of subgraphs upon which the knowledge graph embedding models are trained also needs to be diligently selected. The more the agreement between ranking of the triples in multiple subgraphs; the higher is the ranking of those triples while aggregating the link prediction results across all those subgraphs.

The embedding dimension size is another crucial hyperparameter in our model. The goal that our model seeks to accomplish is to reduce dimensionality of the embeddings across these subgraphs in order to produce good vector representations and make the computation of the node and edge embeddings feasible. Embedding dimension size will be a function of the size of individual subgraphs as it is directly proportional to the size of individual subgraphs sampled from the KG.

Equations 2, 3, 4 and 5 showcase the effect of hyperparameters relevant to the optimization of the ensemble learner:

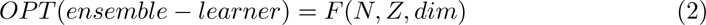

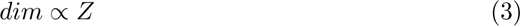

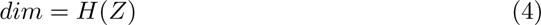

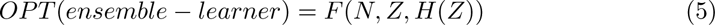

where OP T (*ensemble learner*) denotes the optimization of the ensemble learner as a function *F* () of the number of sampled subgraphs *N*, size of individual subgraphs *Z*and embedding dimension size *dim* (hyperparamter for the individual subgraph-learner) which in turn is a function *H*() of the size of individual subgraphs Z i.e. *H*(*Z*).

### 3.4 Evaluating the performance on link prediction tasks from each individual subgraph

Evaluation of link predictions from individual subgraphs is performed on the test set which is created prior to the graph sampling process. Evaluation metrics include *hits*@*k*, mean rank *MR* and adjusted mean rank *AMR* [40].

One notable issue with evaluating each subgraph separately is the influence of subgraph sizes on ranking metrics. Previous work has shown that these metrics are dependent on test set sizes [41]. Here, using a toy example, we showcase that ranking metrics are also affected by subgraph sizes.

#### 3.4.1 Calculation of rank-based evaluation metrics

Let us consider a knowledge graph *G* with a set of entities *E*, a set of relations *R*, and an embedding scoring function *F* (). Evaluation of a test triple (*h_E_, r_R_, t_E_*) ∈ *T*, includes both right-side as well as left-side evaluations which are analogous to each other and can be exchanged without loss of generality.

To perform right-side evaluation of (*h, r, t*), a set of candidate triples *C*= {(*h, r, t^′^*) — *t^′^* ∈} and a set of plausibility scores for each candidate triple *S*={*F*(*h, r, t^′^*) — *t^′^* ∈ are generated using the scoring function of the embedding model *F* (*h, r, t^′^*) and to perform left-side evaluation of (*h, r, t*), a set of candidate triples *C*={(*h^′^, r, t*) — *h^′^* ∈} and a set of plausibility scores for each candidate triple *S*={*F* (*h^′^, r, t*) — *h^′^* ∈} are generated using the scoring function of the embedding model *F* (*h^′^, r, t*). We need to keep in mind that the ground-truth triple (*h, r, t*) ∈ *C*. *S* is then sorted in descending order based on the plausibility score allotted to each triple in *S* by the scoring function *F* (*h, r, t*). The ranking of the ground-truth triple i.e. *rank*(*h, r, t*) is computed as the index of *F* (*h, r, t*) in the sorted set *S*. The evaluation metrics are calculated as follows, with |*T*| being the cardinality of the test-set *T*.

Mean rank is calculated using equation 6:

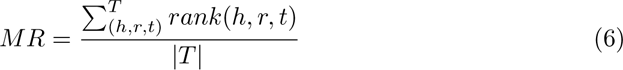

and Hits@k is calculated using equation 7:

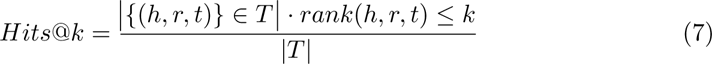

#### 3.4.2 Influence of graph size on evaluation metrics

Here, we show that the rank-based evaluation metrics are influenced by graph size for anon-ideal knowledge graph embedding model. For illustrating the influence of the graph size on rank-based evaluation metrics we use a toy example, where we only consider the right-side evaluation rank of triples and compute two rank based evaluation metrics: *MR* and *hits*@1.

Let us consider a full knowledge graph *G* which has a set of entities *E_G_*={*e*_1_*, e*_2_*, e*_3_*, e*_4_*, e*_5_}. We sample a subgraph *G^′^* from *G* with a smaller set of entities *E_G_′*= *e*_1_*, e*_2_*, e*_3_. In this toy example, we assume that the scoring function *F* () assigns different plausibility scores for every pair of triples and no candidate triple in *C* = {(*h, r, t^′^*) *| t^′^∈ E_G_*} is filtered out during the evaluation process. Assuming we have a large test set *T* for our example; the first element in *T* is (*h, r, t* = *e*_1_), the second element in *T* is (*h, r, t* = *e*_2_), the third element in *T* is (*h, r, t* = *e*_3_) and so on.

For an ideal embedding model, the score function *F* () will always have the property *h, r, t, t^′^G, F* (*h, r, t*) ¿ *F* (*h, r, t^′^*) (*h, r, t*) ∈ *G and* (*h, r, t^′^*) ∉ *G*. Thus, *F* () will always assign a higher plausibility score for a true triple compared to a false triple. Therefore, in both the full graph *G* and subgraph *G^′^*, the rank of a ground-truth triple (*h, r, t* = *e*_1_) is *rank*(*h, r, t* = *e*_1_) = 1. For the full test set *T* ; if this ideality held for every test triple, then the evaluation metrics will be *MR*=1 and *hits*@1 = 1.0 for both *G* and *G^′^*.

However, for a non-ideal embedding model in which the above property does not hold for the scoring function *F* (); we consider the worst case scenario in which *F* () performs no better than a random number generator. In this case, for the full graph *G* with say 5 candidate triples, the probability of a ground-truth triple (*h, r, t* = *e*_1_) to be ranked in the top 1 is 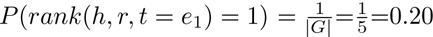. On the other hand, for the subgraph *G*^′^ with only 3 candidate triples, the probability of a ground-truth triple (*h, r, t* = *e*_1_) to be ranked in the top 1 is 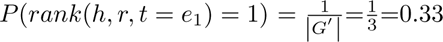. For the large test set *T*, in the case of the full graph *G* with 5 candidate triples the evaluation metrics will converge to the following values that are also shown in Table 2:

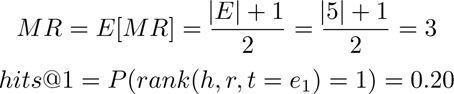

**Table 2.**
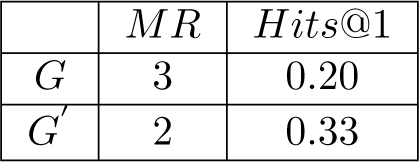
The below table shows the computed mean rank *MR* and *Hits*@1 for our toy example. These are two of the rank-based evaluation metrics that we use to explain the confounding effect of graph size that prevents a fair and direct comparison between these rank-based evaluation metrics achieved on the full KG and our smaller sampled subgraphs.

| | $MR$ | $\text{Hits@1}$ |
| --- | --- | --- |
| $G$ | 3 | 0.20 |
| $G'$ | 2 | 0.33 |

For the large test set *T*, in the case of the subgraph *G^′^* with 3 candidate triples the evaluation metrics will converge to the following values that are also shown in Table 2:

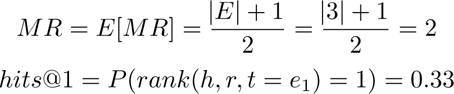

As illustrated by our toy example, for a realistic non-ideal embedding model, smaller graph sizes can yield a lower *MR* and higher *hits*@*k*, thus leading to a bias in our link prediction evaluations. Therefore, this confounder prevents a fair and direct comparison between the rank-based metrics of a full knowledge graph and our smaller sampled subgraphs.

#### 3.4.3 Size-independent evaluation metrics

There are a few evaluation metrics that are less dependent on the size of the graph. Notably, the adjusted mean rank (*AMR*) corrects the *MR* metric according to the graph size [41]:

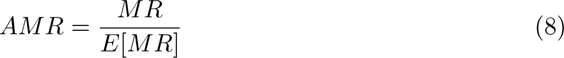

In equation 8 *MR* represents the mean rank of the link predictions, *E*[*MR*] represents the expected mean rank and *AMR* is the adjusted mean rank metric. *AMR* ranges in the interval [0, 2]. In order to make adjusted mean rank metric more intuitive we define another metric called adjusted mean rank index *AMRI* = 1 *AMR* that ranges in the interval [-1,1] where a value closer to 1 indicates good performance on link prediction and a value closer to −1 indicates poor performance on link prediction. *AMRI* is one of the size independent evaluation metrics that we use to evaluate the link predictions yielded by our subgraph-learners.

Another size independent evaluation metric that can be potentially considered for evaluating our approach is a percentile-based metric - hits@k%, which represents the fraction of test triples that appears in the top *k^th^* percentile of link predictions. This concept is very similar to *AMR*, since it corrects the rank-based metric *Hits*@*k* according to the size of the graph.

### 3.5 Aggregating the results from the link prediction across multiple subgraphs

In addition to evaluating the individual subgraph-learners, the link predictions from each subgraph need to be aggregated for generating a consolidated set of link predictions across multiple sampled subgraphs. This paper proposes an ensemble method for aggregation of the plausibility scores for ranked triples / link predictions from subgraphs and explores the effects of ensembling from multiple subgraphs on reduction of variance and improvement in link predictions. The two steps involved in this aggregation strategy are:

1. Score normalization for individual subgraphs
2. Score aggregation across multiple subgraphs

#### 3.5.1 Score normalization for individual subgraphs

Since each subgraph and the underlying scoring function *F* () for it’s corresponding learner are trained separately; for a given triple (*h, r, t*) in say subgraphs *G*_1_ and *G*_2_, the plausibility scores generated by scoring function *F*_1_(*h, r, t*) for *G*_1_ and scoring function *F*_2_(*h, r, t*) for *G*_2_ are not directly comparable. In this paper, we utilize a normalizer *norm*() to standardize the plausibility scores assigned by the respective scoring functions of the subgraph-learners and we have explored two normalizers for the same purpose:

- *StandardScaler*(): rescale plausibility scores to mean of 0 and standard deviation of 1.
- *MinMaxScaler*(): rescale plausibility scores to minimum of 0 and maximum of 1.

Both normalizers require the distribution of the plausibility scores in each subgraph. Since knowledge graphs are usually very large, even though we have reduced the graph size by our stochastic sampling approach, each subgraph still contains a large number of possible triple combinations, of the order of *O*( *R E* ) where *R* represents the number of relations in a given subgraph and *E* represents the number of entities in the same subgraph. Therefore, we can only sample a portion of the triples for learning the distribution of the plausibility scores.

Instead of performing random sampling, we propose to learn the distribution of the plausibility scores by leveraging the concept of a negative sampler for the knowledge graph [42]. Assuming an effective scoring function *F* () and a closed-world graph, we expect the scores to be normally-distributed with true triples lying in the higher range of the distribution and false/corrupted triples lying in the lower range of the distribution as shown in Figure 5. True triples can be drawn from the training data whereas false triples can be synthesized using a Bernoulli negative sampler [17], which receives a true triple as input and probabilistically corrupts its head or tail entity to yield a corrupted / false triple as the output.

**Figure 5.**
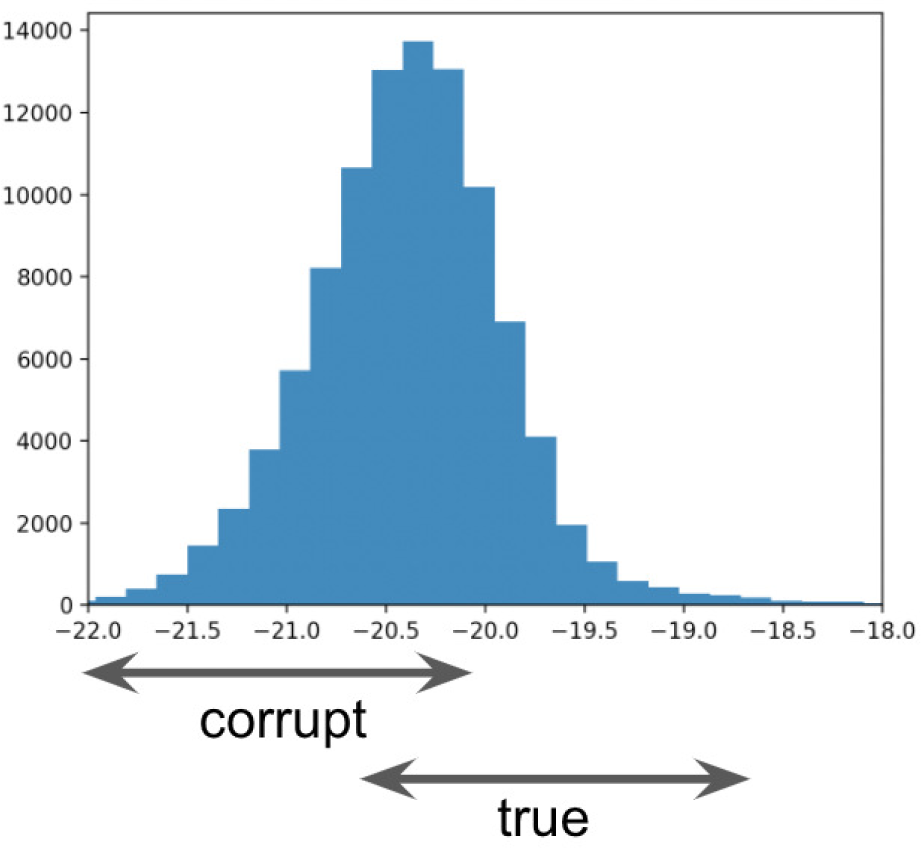
Given a scoring function *F* () and a subgraph; the distribution of scores for the triples in the subgraph *S* is roughly normal where the true triples are in the higher range of the score distribution and the false/corrupted triples are in the lower range of the score distribution.

For each subgraph, a distribution of the scores can then be computed and fitted to agraph-specific *norm*(). During the aggregation phase, scores from each subgraph are transformed using the graph-specific *norm*() before being aggregated into the ensemble model.

#### 3.5.2 Score aggregation across multiple subgraphs

After normalizing the plausibility scores to the same range for the ranked triples / link predictions from each subgraph; the scores need to be aggregated for the full ensemble graph *G_ensemble_* whose set of triples is the same as the original full knowledge graph G, except *F_ensemble_* is an aggregation function *agg*() which combines plausibility scores from all the trained subgraph-learners. In this section, we explore various aggregation functions *agg*() that can be utilized to aggregate normalized plausibility scores from multiple subgraphs to come up with a consolidated set of link predictions for *G_ensemble_*:

- Mean()
- Median()
- Max()

We have used a toy example to illustrate the score aggregation process using two subgraphs *G*_1_ and *G*_2_, assuming ground-truth triple (*h, r, t* = *e*_1_), normalizer *norm*() = *MinMaxScaler*() and aggregation function *agg*() = *Mean*(). The scores shown in Table 3 are for illustration purposes only.

**Table 3.**
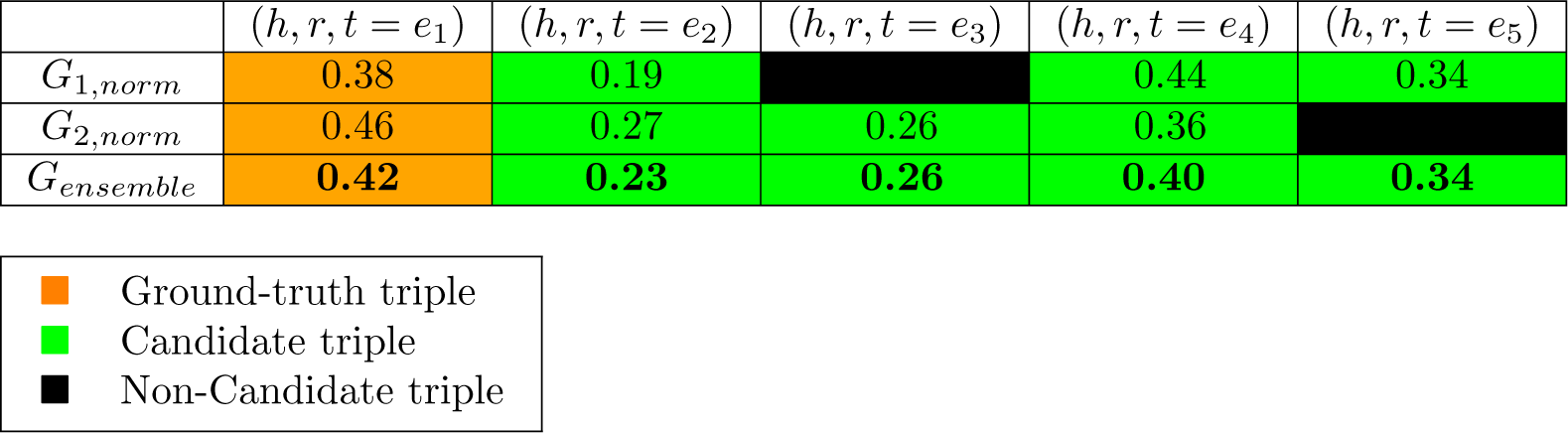
Illustration of the score aggregation process using two toy subgraphs *G*_1_ and *G*_2_. Triple (*h, r, t* = *e*_3_) is not a candidate triple in subgraph *G*_1_ since entity *e*_3_ ∈ *G*_1_. Triple (*h, r, t* = *e*_5_) is not a candidate triple in subgraph *G*_2_ since entity *e*_5_ ∈ *G*_2_. *G_ensemble_* is the *mean*() of the normalized plausibility scores of a triple (*h, r, t*) in *G*_1_*_,norm_* and *G*_2_*_,norm_* assuming that triple (*h, r, t*) ∈ *G*_1_*andG*_2_.

| | $(h, r, t = e_1)$ | $(h, r, t = e_2)$ | $(h, r, t = e_3)$ | $(h, r, t = e_4)$ | $(h, r, t = e_5)$ |
| --- | --- | --- | --- | --- | --- |
| $G_{1,norm}$ | 0.38 | 0.19 | | 0.44 | 0.34 |
| $G_{2,norm}$ | 0.46 | 0.27 | 0.26 | 0.36 | |
| $G_{ensemble}$ | 0.42 | 0.23 | 0.26 | 0.40 | 0.34 |

|  |  |
| --- | --- |
| ■ | Ground-truth triple |
| ■ | Candidate triple |
| ■ | Non-Candidate triple |

After aggregation of the normalized plausibility scores, the ranks of the triples in the test set are computed and standard evaluation metrics such as *MR*, *AMR* and *hits*@*k* are calculated for the consolidated set of link predictions that are generated as a result of this aggregation. The advantage of performing the evaluation on *G_ensemble_* instead of each individual subgraph is that both *G* and *G_ensemble_* are of the same size and therefore *G* and *G_ensemble_* have the same number of entities (*|E_ensemble_|* = *|E|*) and relations ( *R_ensemble_* = *R* ). So; graph size is eliminated as a confounder during evaluation. Therefore, direct comparison of the rank-based metrics between *G* and *G_ensemble_* is fair and warranted.

## 4 Experimental Setup

The knowledge graph embedding models in the following experiments were trained using PyKEEN [43] on a high-performance computing cluster that used a V100 Volta GPU with 32 GB of memory on 4 biomedical KG datasets:

1. Drug Repurposing Knowledge Graph (DRKG) [13]
2. Integrated Biomedical Knowledge Hub (iBKH) [14]
3. Hetionet [11, 12]
4. BioKG [10]

Two types of experiments were conducted and evaluation metrics were computed based on the performance of the ensemble learner in generating relevant link predictions in both of those experiments. The two types of experiments that were conducted have been described below:

1. Task specific link prediction on large biomedical knowledge graph datasets.
2. General link prediction on both large as well as relatively smaller biomedical knowledge graph datasets.

Normalizer used to normalize plausibility scores for ranked triples / link predictions from each subgraph: *MinMaxScaler*()

Aggregator function used for aggregating the plausibility scores for triples across multiple subgraphs: *Mean*()

Hyperparameter optimization:

- Embedding dimensions: [3000, 4000, 5000, 6000]
- Number of negatives per positive triple: [1, 10, 100]
- Margin for Margin Ranking Loss: [1, 4, 7, 10]
- Regularization coefficients: [0.02, 0.06, 0.10]

Model: RotatE [7] - This embedding model was used to demonstrate the performance boost obtained by ensembling multiple individual instances of subgraph learners that use RotatE to generate embeddings in the localized view space of the subgraphs. The reason for selecting this embedding model over the other KGE models is because it outperformed all the other KGE models on link prediction when they were trained on the full scope of the KG on all the four biomedical graph datasets. Further details regarding the reason for the selection of this embedding model for training the individual subgraph learners have been mentioned in Table 4 under the Results and Discussion section of this article.

**Table 4.** Evaluation metrics obtained by various knowledge graph embedding models on link prediction when trained on the full scope of the KG for the four benchmark biomedical knowledge graph datasets namely Bio-KG, Hetionet, DRKG and iBKH.

| TransE |  |  |  |  |
| --- | --- | --- | --- | --- |
| KG | *MR* | *AMRI* | *Hits@1* | *Hits@10* |
| Bio-KG | 712.2 | 0.96 | 0.019 | 0.24 |
| Hetionet | 667.6 | 0.97 | 0.034 | 0.26 |
| DRKG | 929.4 | 0.84 | 0.023 | 0.19 |
| iBKH | 3468.2 | 0.59 | 0.007 | 0.11 |
| RotatE |  |  |  |  |
| KG | *MR* | *AMRI* | *Hits@1* | *Hits@10* |
| Bio-KG | 634.2 | 0.976 | 0.059 | 0.28 |
| Hetionet | 413.6 | 0.998 | 0.063 | 0.33 |
| DRKG | 463.9 | 0.99 | 0.135 | 0.31 |
| iBKH | 1480.2 | 0.785 | 0.068 | 0.147 |
| ComplEx |  |  |  |  |
| KG | *MR* | *AMRI* | *Hits@1* | *Hits@10* |
| Bio-KG | 5226.7 | -0.102 | 0.003 | 0.011 |
| Hetionet | 4235.6 | 0.44 | 0.01 | 0.10 |
| DRKG | 684.2 | 0.972 | 0.03 | 0.26 |
| iBKH | 4982.8 | 0.41 | 0.008 | 0.092 |
| DistMult |  |  |  |  |
| KG | *MR* | *AMRI* | *Hits@1* | *Hits@10* |
| Bio-KG | 3572.2 | 0.198 | 0.007 | 0.036 |
| Hetionet | 5184.2 | 0.278 | 0.011 | 0.088 |
| DRKG | 24032.3 | -0.116 | 0.000 | 0.012 |
| iBKH | 312651.8 | -0.388 | 0.000 | 0.005 |

To avoid overfitting the knowledge graph embedding models to any of the individual subgraphs we create a holdout validation set of triples and the validation loss that is obtained on this set is used for early-stopping of the algorithm.

## 5 Results and Discussion

We conducted several experiments to observe the performance of our ensemble learner for link prediction on biomedical knowledge graph datasets and generated evaluation metrics to compare and quantify the performance of our ensemble learner with that of the traditional approach of training KGE models on full scope of the knowledge graph.

Firstly we trained standard knowledge graph embedding models on the full KG for all the four benchmark biomedical knowledge graph datasets namely Bio-KG, Hetionet, DRKG and iBKH to generate entity and relation embeddings. Then we evaluated the performance of these embeddings on link prediction tasks in the test set. Thereafter we computed evaluation metrics such as Mean rank *MR*, Arithmetic Mean rank index *AMRI*, *Hits*@3 and *Hits*@10 to understand which knowledge graph embedding model performed the best on the link prediction tasks across the four biomedical KG datasets that we considered. We used this experiment as the basis to select the KGE model that we used for training the individual subgraphs in our architecture and to further demonstrate the performance boost upon ensembling the individual subgraph learners that are trained using the very same embedding model.

As shown in Table 4; we have computed four different evaluation metrics for the link predictions generated by the standard KGE models on the full benchmark biomedical knowledge graph datasets namely:

1. Mean Rank (*MR*)
2. Adjusted mean rank index (*AMRI*)
3. *Hits*@1
4. *Hits*@10

As shown in Table 4 RotatE undisputedly outperformed all the other KGE models for link prediction on all the four benchmark biomedical KG datasets that we have considered. Therefore; as a result we selected RotatE to train our individual subgraph learners prior to ensembling the link predictions. We have demonstrated the performance boost that is achieved upon ensembling the link predictions from individual subgraph learners over generating link predictions using a KGE model trained on the full KG.

The evaluation results of our ensemble learner have been reported for general link prediction on all the four benchmark biomedical KG datasets regardless of their size and for task-specific (drug-disease) link prediction on the relatively larger biomedical KG datasets among the four benchmark datasets namely DRKG and iBKH.

The various experiments that we conducted for evaluating our ensemble learner have been listed below:

1. DRKG : Compound → Disease task specific link prediction
2. iBKH : Drug → Disease task specific link prediction
3. DRKG : General link prediction
4. iBKH : General link prediction
5. Hetionet : General link prediction
6. Bio-KG : General link prediction

### 5.1 DRKG : Compound → Disease task specific link prediction

For the task-specific (drug-disease) link prediction experiment, subgraphs were sampled from the full DRKG knowledge graph as shown in Figure 3 and all the entities belonging to the drug and disease categories were added into the sampled subgraphs so that each subgraph-learner had all the context that it needed to come up with good link predictions for the (drug-disease) entity pairs. To start with we sampled 3 subgraphs from the full knowledge graph *G*. Structure and metadata of the full knowledge graph *G* as well as the 3 sampled subgraphs *G*_1_, *G*_2_ and *G*_3_ are shown in Table 5. Each subgraph contains 3M triples in the training set and 350k triples in the validation set. The test set is the same across all the subgraphs and only contains 7,857 triples since only triples of the type (drug, relation, disease) are kept and the rest are filtered out of the test set.

**Table 5.** Structure of the full DRKG graph and the 3 subgraphs sampled from it. This example is to simply show how the triples are distributed per sampled subgraph. We generally sample more number of subgraphs before ensembling the link predictions from models trained on each of them so as to reduce the variance and obtain link predictions from more localized view spaces.

|  | Entities | Relations | Training | Validation | Testing |
| --- | --- | --- | --- | --- | --- |
| $G$ | 97,238 | 107 | 5,800k | 500k | 7,857 |
| $G_1$ | 36,440 | 97 | 3,127k | 348k | 7,857 |
| $G_2$ | 36,390 | 92 | 3,043k | 338k | 7,857 |
| $G_3$ | 37,760 | 95 | 3,187k | 354k | 7,857 |
| $G_{ensemble}$ | 97,238 | 107 | 3,800k | 500k | 7,857 |

The evaluation metrics for this experiment have been shown in Table 6 for the RotatE model trained on the full knowledge graph *G* as well as the individual subgraphs *G*_1_,*G*_2_, *G*_3_ that were sampled from *G* and for the ensemble learner on the aggregated graph *G_ensemble_*. The model trained on the full knowledge graph *G* demonstrated better mean rank *MR* and adjusted mean rank index *AMRI* for the task-specific (drug, disease) link predictions compared to the model trained on the individual subgraphs *G*_1_,*G*_2_, *G*_3_ that were sampled from *G* but their respective *Hits*@*k* were almost similar. On the other hand, the consolidated task-specific (drug, disease) link predictions *G_ensemble_* showed better mean rank *MR*, adjusted mean rank index *AMRI* and *Hits*@*k* compared to the link predictions generated by the models trained on the individual subgraphs *G*_1_,*G*_2_ and *G*_3_ as well as the full knowledge graph *G*.

**Table 6.** Results of ensembling link predictions from 3 subgraphs and aggregating their respective plausibility scores. *G_ensemble_* is aggregated from 3 subgraphs *G*_1_, *G*_2_ and *G*_3_. Link predictions obtained from *G_ensemble_* perform better than the link predictions obtained from the full DRKG knowledge graph *G*.

| | $MR$ | $AMRI$ | $Hits@1$ | $Hits@3$ | $Hits@10$ |
| --- | --- | --- | --- | --- | --- |
| $G$ | **746.1** | **0.985** | **0.034** | **0.075** | **0.154** |
| $G_1$ | 848.6 | 0.953 | 0.037 | 0.079 | 0.159 |
| $G_2$ | 939.3 | 0.948 | 0.035 | 0.075 | 0.160 |
| $G_3$ | 953.9 | 0.949 | 0.035 | 0.077 | 0.162 |
| $G_{ensemble}$ | **668.9** | **0.986** | **0.045** | **0.092** | **0.188** |

Instead of only ensembling link predictions from 3 subgraphs, we adjust the hyperparameter controlling the number of sampled subgraphs in order to show our evaluation results on ensembling link predictions from 10 subgraphs *G*_1_ − *> G*_10_ in Table 7. This was primarily done to reduce the variance and aggregate plausibility scores from more number of localized view spaces. The results observed with 10 subgraphs were similar to the trend observed with 3 subgraphs, in which link predictions from the subgraphs yielded similar *Hits*@*k* compared to the link predictions from the full graph *G* but the consolidated link predictions yielded by the ensemble learner on *G_ensemble_* performed better in all the evaluation metrics compared to the KGE model trained on the full knowledge graph *G*.

**Table 7.** Results of ensembling link predictions from 10 subgraphs and aggregating their respective plausibility scores.Link predictions from *G_ensemble_* (which is aggregated from 10 subgraphs *G*_1_→ *G*_10_) performs better than the link predictions from the full DRKG graph *G*. The test set only includes (Drug, Disease) triples.

|  | MR | AMRI | Hits@1 | Hits@3 | Hits@10 |
| --- | --- | --- | --- | --- | --- |
| $G$ | **746.1** | **0.985** | **0.034** | **0.075** | **0.154** |
| $Mean(G)$ | 937.9 | 0.949 | 0.036 | 0.076 | 0.157 |
| $Std(G)$ | (32.3) | (0.002) | (0.001) | (0.001) | (0.003) |
| $G_{ensemble}$ | **606.7** | **0.987** | **0.050** | **0.101** | **0.195** |

### 5.2 iBKH : Drug → Disease task specific link prediction

For the task-specific (drug-disease) link prediction experiment, subgraphs were sampled from the large iBKH knowledge graph as shown in Figure 3 and all the entities belonging to the drug and disease categories were added into the sampled subgraphs so that each subgraph-learner had all the context that it needed to come up with good link predictions for the (drug-disease) entity pairs. Compared to the 6M triples in DRKG, iBKH is much larger with 50M triples. Therefore, when we train a RotatE model to generate embeddings for the full iBKH graph; even at the maximum embedding dimension size of 6000, the performance on task-specific (drug-disease) link prediction remains poor as shown by the evaluation metrics in (Table 8). It is expected that increasing the embedding dimension sizes further might improve the link prediction capabilities on this massive KG. However; using an embedding dimension size of 6000 to represent the graph data was already enough to max out our GPU memory. So, clearly using a single KGE model to generate vector representations for the full KG in this case is infeasible due to the massive size of the iBKH KG. Therefore we sample 10 subgraphs *G*_1_ *> G*_10_ from the full KG *G* using our stochastic sampling approach to conduct this experiment on iBKH. We ensemble link predictions from more number of subgraphs in order to reduce the variance and aggregate plausibility scores from multiple localized view spaces. We also summarized the evaluation metrics obtained across all the sampled subgraphs by computing the Mean *Mean*(*G*) and Standard deviation *Std*(*G*) statistic for each of them. The results that we achieved followed the same trend that we observed when ensembling task-specific link predictions from multiple subgraphs for DRKG in which the consolidated set of task-specific link predictions yielded by the ensemble learner on *G_ensemble_* performed much better in all the evaluation metrics compared to the KGE model trained on the full knowledge graph *G*.

**Table 8.**
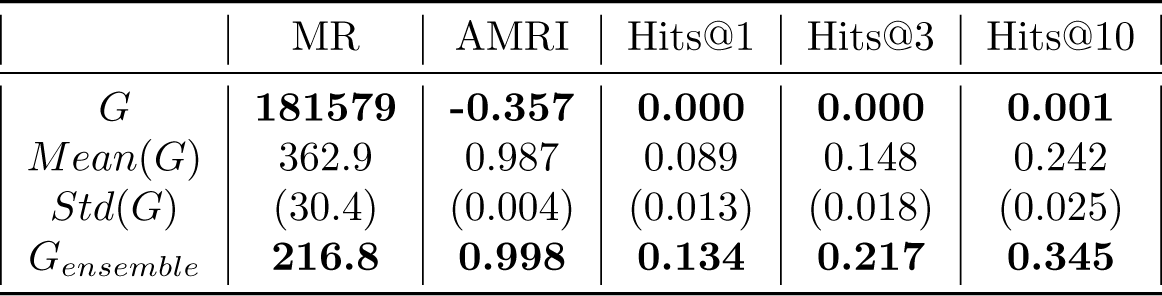
Results of ensembling link predictions from 10 subgraphs and aggregating their respective plausibility scores for iBKH. Link predictions from *G_ensemble_* (which is aggregated from 10 subgraphs *G*_1_→ *G*_10_) performs better than the link predictions from the full iBKH graph *G*. The test set included only includes (Drug, Disease) triples.

Based on the evaluation metrics shown in Table 8; we concluded that dividing the full iBKH graph into multiple smaller subgraphs and aggregating the individual subgraph learners / embedding models on *G_ensemble_* demonstrated significantly improved performance compared to the KGE model that was trained on the full iBKH graph *G*.

### 5.3 DRKG : General link prediction

General link prediction experiment was conducted on DRKG to demonstrate our ensemble learner’s capabilities to yield good link predictions between any two types of entity pairs from the KG on a large biomedical KG. This experiment was conducted in addition to the task-specific (drug-disease) link prediction experiment shown previously in this section for DRKG because showcasing that our model can generalize well to predict links between any two categories of entities in the KG is crucial to truly understand if the performance boost previously observed in the task-specific link prediction experiment was indeed a capability of our ensemble learner or some inherent bias in the KG topology relevant to the (drug, disease) entity pairs. Table 10 shows the evaluation metrics on the consolidated set of link predictions aggregated from 20 subgraphs by our ensemble-learner for iBKH KG. The *Hits*@10 metric for the link predictions yielded by our ensemble learner trained on the group of 20 sampled subgraphs *G_ensemble_* showed a 34% increase compared to the *Hits*@10 metric for the link predictions yielded by an individual KGE model trained on the full knowledge graph *G*.

Table 9 shows the evaluation metrics on the consolidated set of link predictions aggregated from 10 subgraphs for DRKG; even though mean rank *MR* is lower in *G_ensemble_* compared to full KG *G*; *hits*@10 showed a 10% improvement compared to the individual KGE model trained on *G*. We have also plotted the variation of the evaluation metrics (*Hits*@1, *Hits*@3, *Hits*@10, *MR* and *AMRI*) for the consolidated set of link predictions against the number of subgraphs used for aggregating those link predictions in DRKG as shown in Figure 6.

**Figure 6.**
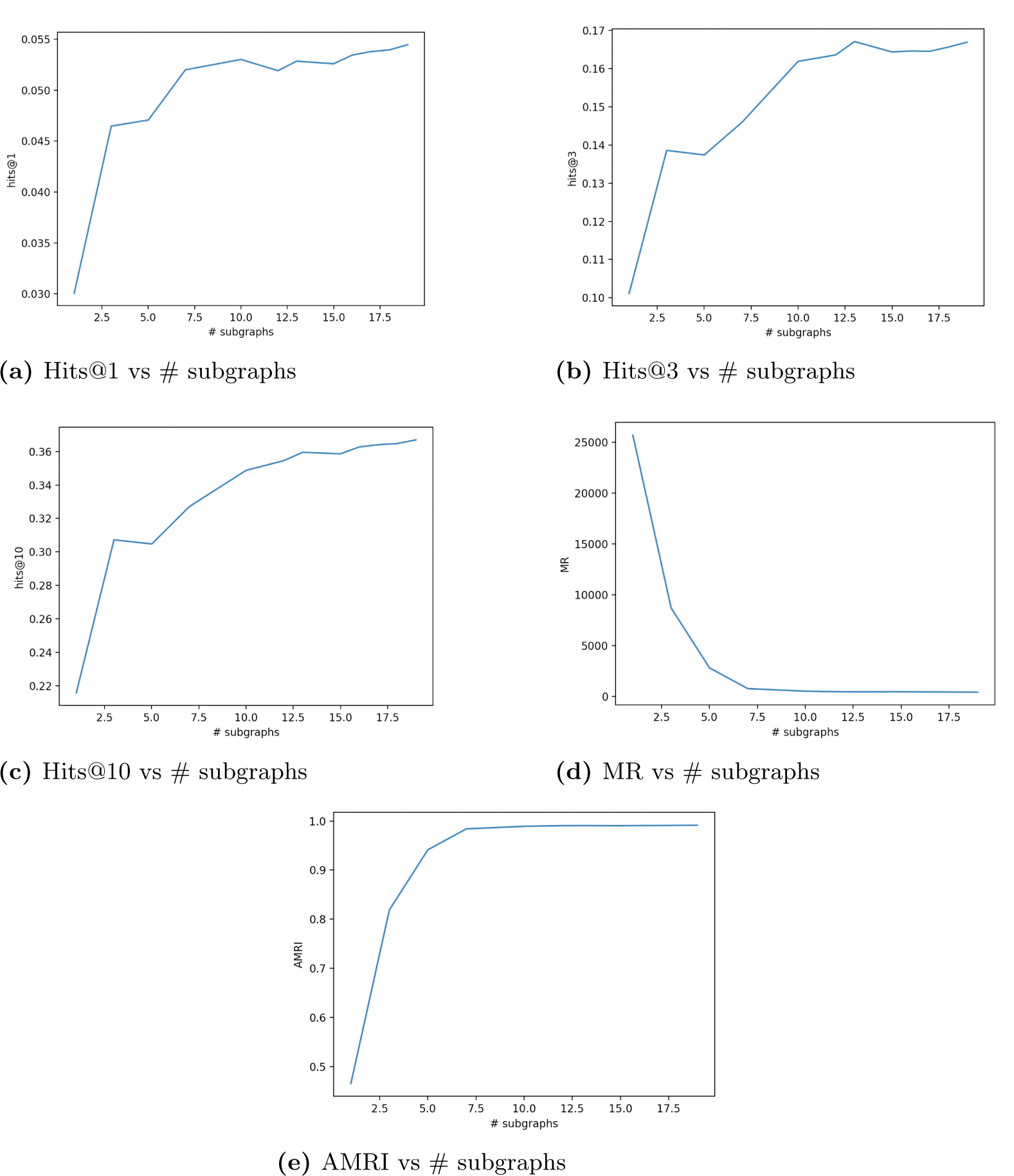
Plots visualizing the variation of evaluation metrics for the consolidated set of link predictions in *G_ensemble_* with the number (#) of subgraphs used for consolidating / aggregating those link predictions in the case of DRKG. The plots include : (a) Hits@1 vs number of subgraphs, (b) Hits@3 vs number of subgraphs, (c) Hits@10 vs number of subgraphs, (d) MR vs number of subgraphs, (e) AMRI vs number of subgraphs.

**Table 9.** Result of ensembling link predictions from 10 sampled subgraphs in the case of DRKG. The test set included all types of triples as this was a general link prediction experiment.

|  | MR | AMRI | Hits@1 | Hits@3 | Hits@10 |
| --- | --- | --- | --- | --- | --- |
| $G$ | **463.9** | **0.990** | **0.044** | **0.135** | **0.311** |
| $Mean(G)$ | 264.6 | 0.982 | 0.193 | 0.442 | 0.601 |
| $Std(G)$ | (93.8) | (0.013) | (0.057) | (0.033) | (0.038) |
| $G_{ensemble}$ | **521.0** | **0.989** | **0.053** | **0.162** | **0.349** |

### 5.4 iBKH : General link prediction

General link prediction experiment was conducted on iBKH to demonstrate our ensemble learner’s capabilities to yield good link predictions between any two types of entity pairs from the KG on a large biomedical KG. This experiment was conducted in addition to the task-specific (drug-disease) link prediction task shown previously in this section for iBKH because showcasing that our model can generalize well to predict links between any two categories of entities in the KG is crucial to truly understand if the performance boost previously observed in the task-specific link prediction experiment was indeed a capability of our approach or some inherent bias in the KG topology relevant to the (drug, disease) entity pairs. Since iBKH is a very large knowledge graph with about 50M triples we sampled more number of subgraphs to reduce the variance and ensure that each individual subgraph size was not too large in itself. Table 10 shows the evaluation metrics on the consolidated set of link predictions aggregated from 20 subgraphs by our ensemble-learner for iBKH KG. The *Hits*@10 metric for the link predictions yielded by our ensemble learner trained on the group of 20 sampled subgraphs *G_ensemble_* increased by 0.051 points compared to the *Hits*@10 metric for the link predictions yielded by an individual KGE model trained on the full knowledge graph *G*.

**Table 10.** Result of ensembling link predictions from 20 sampled subgraphs in the case of iBKH. The test set included all types of triples as this was a general link prediction experiment.

|  | MR | AMRI | Hits@1 | Hits@3 | Hits@10 |
| --- | --- | --- | --- | --- | --- |
| $G$ | **1480.2** | **0.785** | **0.007** | **0.068** | **0.147** |
| $Mean(G)$ | 412.27 | 0.956 | 0.202 | 0.182 | 0.32 |
| $Std(G)$ | (88.84) | (0.011) | (0.032) | (0.004) | (0.024) |
| $G_{ensemble}$ | **1240.4** | **0.827** | **0.01** | **0.072** | **0.198** |

Dividing the full iBKH graph into multiple smaller subgraphs and aggregating the link predictions from multiple subgraph learners, *G_ensemble_* demonstrated significantly improved performance on all the evaluation metrics compared to the link predictions generated by training a KGE model on the full graph. We have also plotted the variation of the evaluation metrics (*Hits*@1, *Hits*@3, *Hits*@10, *MR* and *AMRI*) for the consolidated set of link predictions against the number of subgraphs used for aggregating those link predictions in iBKH as shown in Figure 7.

**Figure 7.**
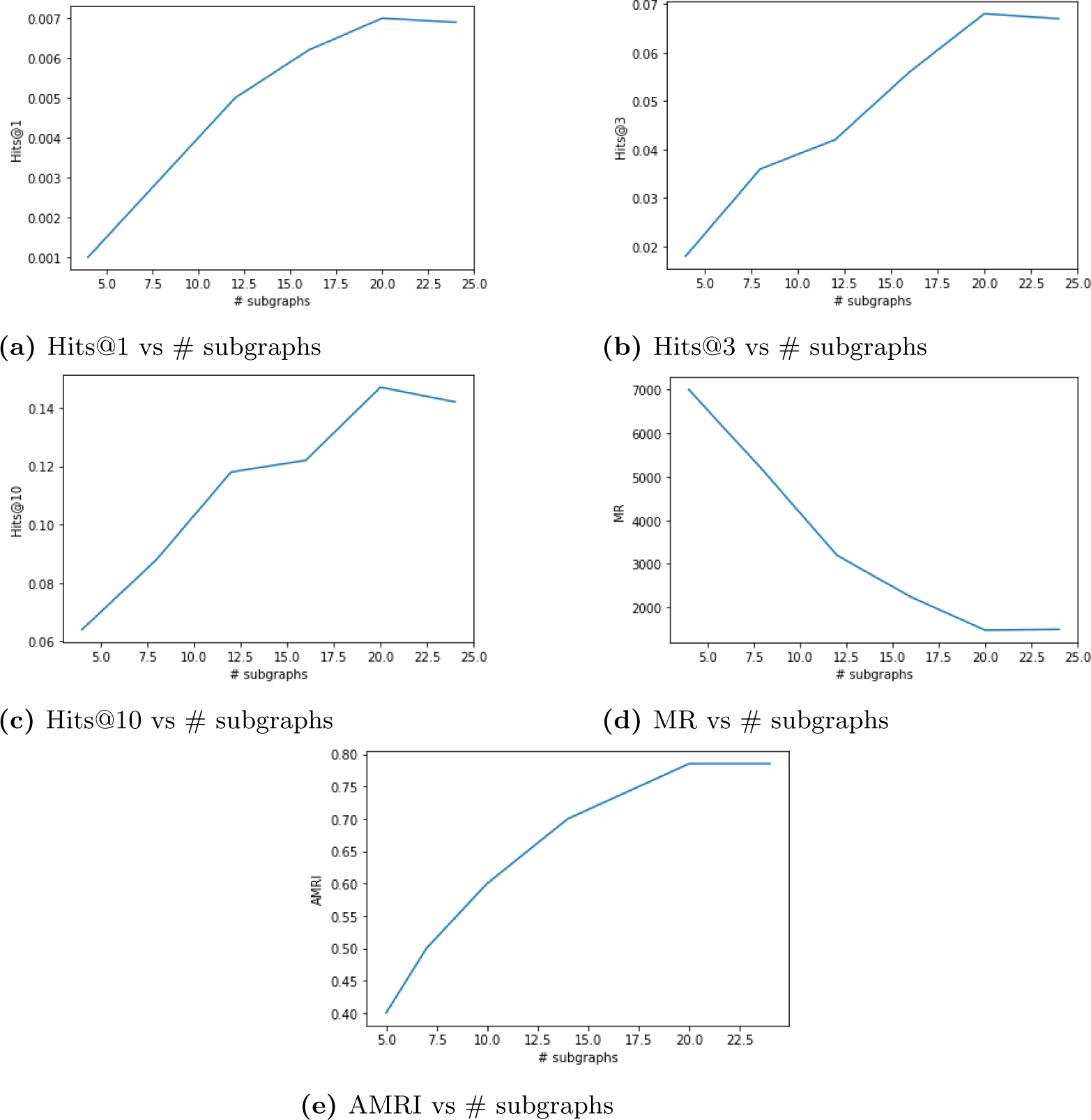
Plots visualizing the variation of evaluation metrics for the consolidated set of link predictions in *G_ensemble_* with the number (#) of subgraphs used for consolidating / aggregating those link predictions in the case of iBKH. The plots include : (a) Hits@1 vs number of subgraphs, (b) Hits@3 vs number of subgraphs, (c) Hits@10 vs number of subgraphs, (d) MR vs number of subgraphs, (e) AMRI vs number of subgraphs.

### 5.5 Hetionet : General link prediction

General link prediction experiment was conducted on Hetionet to demonstrate our ensemble learner’s capabilities to yield good link predictions between any two types of entity pairs from the KG. Table 11 shows the evaluation metrics upon ensembling link predictions from 10 subgraphs to create a consolidated set of link predictions for Hetionet graph dataset. For a relatively smaller KG such as hetionet, the evaluation metrics did not get boosted upon ensembling link predictions from multiple subgraphs. The metrics show that the performance of a KGE model trained on the full graph is almost the same as the performance of an ensemble learner that consolidates link predictions from multiple subgraphs. No significant boost in evaluation metrics was observed between full graph *G* and aggregated graph *G_ensemble_* for Hetionet. We have also plotted the variation of the evaluation metrics (*Hits*@1, *Hits*@3, *Hits*@10, *MR* and *AMRI*) for the consolidated set of link predictions against the number of subgraphs used for aggregating those link predictions in Hetionet as shown in Figure 8.

**Figure 8.**
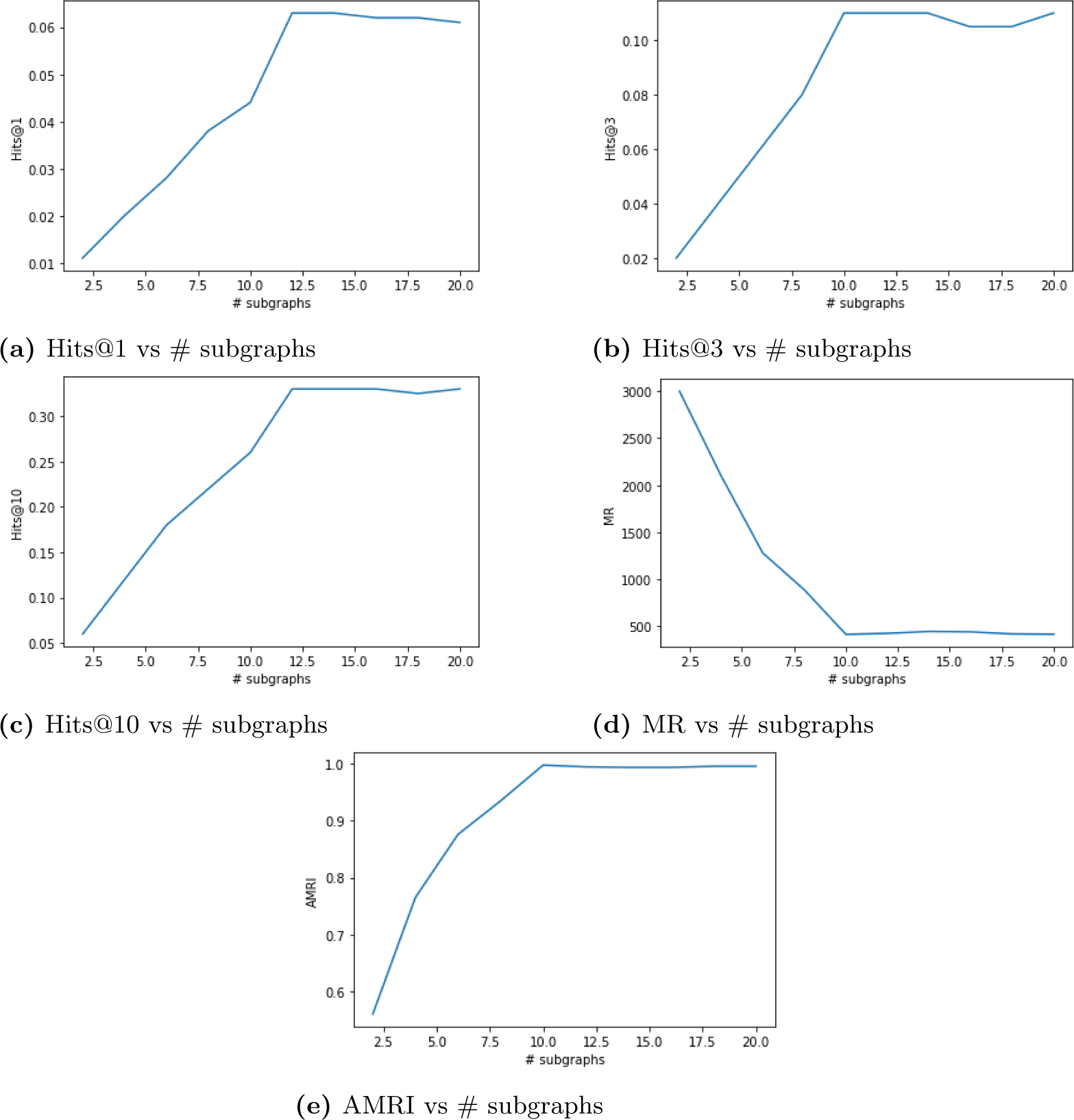
Plots visualizing the variation of evaluation metrics for the consolidated set of link predictions in *G_ensemble_* with the number (#) of subgraphs used for consolidating /aggregating those link predictions in the case of Hetionet. The plots include : (a) Hits@1 vs number of subgraphs, (b) Hits@3 vs number of subgraphs, (c) Hits@10 vs number of subgraphs, (d) MR vs number of subgraphs, (e) AMRI vs number of subgraphs.

**Table 11.** Result of ensembling link predictions from 10 sampled subgraphs in the case of Hetionet. The test set included all types of triples as this was a general link prediction experiment.

|  | MR | AMRI | Hits@1 | Hits@3 | Hits@10 |
| --- | --- | --- | --- | --- | --- |
| $G$ | **413.6** | **0.998** | **0.063** | **0.11** | **0.33** |
| $Mean(G)$ | 224.3 | 0.980 | 0.182 | 0.24 | 0.59 |
| $Std(G)$ | (66.64) | (0.017) | (0.044) | (0.02) | (0.01) |
| $G_{ensemble}$ | **438.2** | **0.994** | **0.062** | **0.107** | **0.328** |

### 5.6 Bio-KG : General link prediction

General link prediction experiment was conducted on Bio-KG to demonstrate our ensemble learner’s capabilities to yield good link predictions between any two types of entity pairs from the KG. Table 12 shows the evaluation metrics upon ensembling link predictions from 10 subgraphs to create a consolidated set of link predictions for Bio-KG dataset. For a relatively smaller graph such as Bio-KG; the evaluation metrics did not get boosted upon ensembling link predictions from multiple subgraphs. In fact, the RotatE model trained on the full knowledge graph seems to be performing better by a very small margin. The metrics show that the performance of a KGE model trained on the full graph to generate link predictions is almost equivalent to the performance of an ensemble learner that aggregates link predictions from multiple subgraphs. No significant boost in evaluation metrics was observed between the full graph *G* and aggregated graph *G_ensemble_*. We have also plotted the variation of the evaluation metrics (*Hits*@1, *Hits*@3, *Hits*@10, *MR* and *AMRI*) for the consolidated set of link predictions against the number of subgraphs used for aggregating those link predictions in Bio-KG as shown in Figure 9.

**Figure 9.**
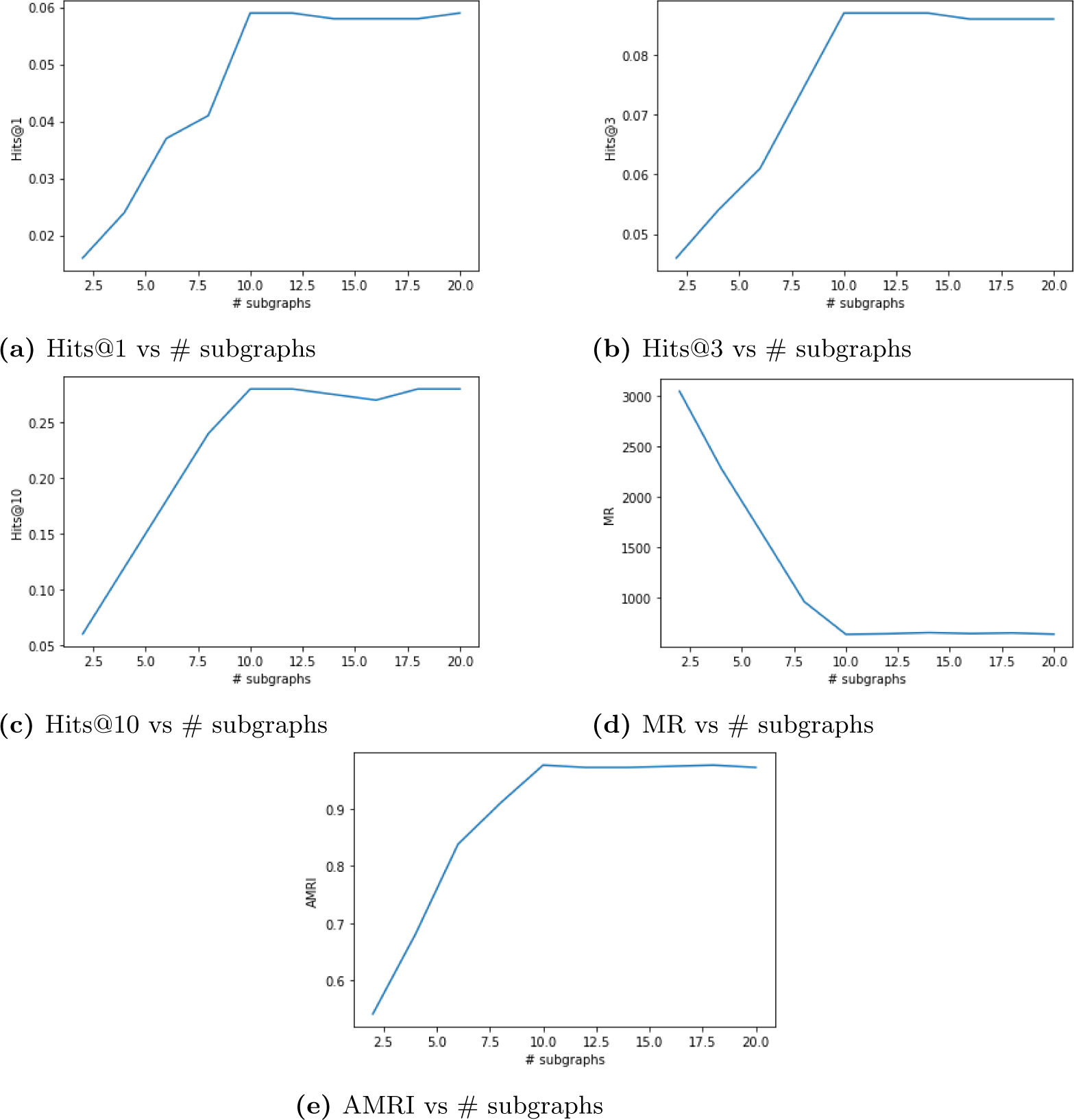
Plots visualizing the variation of evaluation metrics for the consolidated set of link predictions in *G_ensemble_* with the number (#) of subgraphs used for consolidating / aggregating those link predictions in the case of Bio-KG. The plots include : (a) Hits@1 vs number of subgraphs, (b) Hits@3 vs number of subgraphs, (c) Hits@10 vs number of subgraphs, (d) MR vs number of subgraphs, (e) AMRI vs number of subgraphs.

**Table 12.** Result of ensembling link predictions from 10 sampled subgraphs in the case of Bio-KG. The test set included all types of triples as this was a general link prediction experiment.

|  | MR | AMRI | Hits@1 | Hits@3 | Hits@10 |
| --- | --- | --- | --- | --- | --- |
| $G$ | **634.2** | **0.976** | **0.059** | **0.087** | **0.28** |
| $Mean(G)$ | 208.21 | 0.994 | 0.14 | 0.192 | 0.46 |
| $Std(G)$ | (82.2) | (0.02) | (0.003) | (0.01) | (0.02) |
| $G_{ensemble}$ | **667.6** | **0.972** | **0.056** | **0.080** | **0.267** |

Based on the results generated in the above set of experiments it is evident that our ensemble approach which leverages the sampled subgraphs to generate link predictions for the full scope of the knowledge graph outperforms a KGE model that is traditionally trained on the full scope of the knowledge graph in order to generate link predictions for the same. This is because the local views of the full knowledge graph (subgraphs) have lesser noise (unnecessary information that can confuse the KGE model) than the global view. In other words; the KGE models only require a sample of the full KG to predict missing links between a pair of given entities in the KG. This is the underlying concept behind the success of our ensemble learner approach over the traditional approach and it is further validated by the significantly better performance of our ensemble learner in the experiments where we generated link predictions on large biomedical KGs. Additionally since we train the KGE model on smaller sized subgraphs instead of the large knowledge graph; we are also able to reduce the GPU memory footprint associated with training the KGE model.

## 6 Conclusion

In this article, we explored the potential of using a divide and conquer approach to perform knowledge graph completion / link prediction. Specifically, we first performed stochastic sampling of a large knowledge graph into multiple smaller-sized subgraphs using breadth-first search (BFS) on randomly selected seed nodes. Then, we trained knowledge graph embedding models on each individual subgraph to generate embeddings / vector representations in localized view space which were in turn downstreamed to compute plausibility scores for KG triples. We subsequently normalized and aggregated the plausibility score for each triple across multiple subgraphs to generate the consolidated set of link predictions for the full knowledge graph. Experimental results proved that our approach outperforms the traditional approach of training a KGE model on the full graph for link prediction in the case of large knowledge graphs and performs at least as well as the traditional approach of training a KGE model on the full graph for link prediction in the case of smaller knowledge graphs.

## Notes

https://het.io/

https://github.com/gnn4dr/DRKG

https://github.com/wcm-wanglab/iBKH

https://github.com/dsi-bdi/biokg

### Competing Interest Statement

The authors have declared no competing interest.

